# Replicative *Acinetobacter baumannii* strains interfere with phagosomal maturation by modulating the vacuolar pH

**DOI:** 10.1101/2023.02.02.526753

**Authors:** Jesus S. Distel, Gisela Di Venanzio, Joseph J. Mackel, David A Rosen, Mario F. Feldman

**Author notes:** Corresponding author: Mario F. Feldman.

## Abstract

Bacterial pneumonia is a common infection of the lower respiratory tract that can afflict patients of all ages. Multidrug-resistant strains of *Acinetobacter baumannii* are increasingly responsible for causing nosocomial pneumonias, thus posing an urgent threat. Alveolar macrophages play a critical role in overcoming respiratory infections caused by this pathogen. Recently, we and others have shown that new clinical isolates of *A. baumannii*, but not the common lab strain ATCC 19606 (19606), can persist and replicate in macrophages within spacious vacuoles that we called *<u>A</u>cinetobacter* <u>C</u>ontaining <u>V</u>acuoles (ACV). In this work, we demonstrate that the modern *A. baumannii* clinical isolate 398, but not the lab strain 19606, can infect alveolar macrophages and produce ACVs *in vivo* in a murine pneumonia model. Both strains initially interact with the alveolar macrophage endocytic pathway, as indicated by EEA1 and LAMP1 markers; however, the fate of these strains diverges at a later stage. While 19606 is eliminated in an autophagy pathway, 398 replicates in ACVs and are not degraded. We show that 398 reverts the natural acidification of the phagosome by secreting large amounts of ammonia, a by-product of amino acid catabolism. We propose that this ability to survive within macrophages may be critical for the persistence of clinical *A. baumannii* isolates in the lung during a respiratory infection.

## Introduction

The opportunistic bacterial pathogen *Acinetobacter baumannii* is an urgent global public health threat. This pathogen is associated with a wide variety of nosocomial infections including pneumonia, meningitis, urinary tract infection, and wound infection. However, respiratory tract infections are the principal disease associated with *A. baumannii* (Di Venanzio et al., 2019). Alveolar macrophages (AMs) and lung epithelial cells are the first line of defense against respiratory pathogens (Bissonnette et al., 2020; Mubarak et al., 2018).

Macrophages are particularly important during the early stages of *A. baumannii* respiratory infection, as they initiate and regulate the innate immune response against *A. baumannii* by recruiting non-resident macrophages and neutrophils (García-Patiño et al., 2017; Pires & Parker, 2019; H. Qiu et al., 2012). Consequently, depletion of AMs during *A. baumannii* infection results in an increased in bacterial burden, tissue damage, and sepsis in the host (H. H. Lee et al., 2020). Moreover, it has been demonstrated that murine AMs can internalize *A. baumannii in vivo* at 4 h post intranasal infection (H. Qiu et al., 2012).

Phagocytosis is a mechanism by which cells internalize large particles, such as apoptotic cells or bacteria. Phagocytosis is required to maintain cellular homeostasis and is essential in the innate immune response against pathogens (Rosales & Uribe-Querol, 2017; Uribe-Querol & Rosales, 2017). This process is performed primarily by myeloid cells including macrophages, neutrophils, monocytes, and dendritic cells (Lim et al., 2017; Rabinovitch, 1995). After particle recognition and internalization, the newly formed phagosome undergoes a dynamic series of steps called “maturation”. Each phase of the process is determined by the presence of specific signaling molecules located on the phagosomal membrane. Initially, the phagosome recruits the characteristic endosomal marker Rab5 with its effectors EEA1 and VPS34. Maturation continues with the exchange of these markers by late endosome-associated molecules, such as Rab7, HOPS complex, LAMP1 and LAMP2 among others (Fountain et al., 2021; Kinchen & Ravichandran, 2008; H.-J. Lee et al., 2020). The process concludes with the formation of the phagolysosome and the degradation of its internal content by the activity of proteases, nucleases and lipases (Jeschke & Haas, 2018; Nguyen & Yates, 2021; Westman et al., 2020). During maturation phagosomes progressively decrease luminal pH, mainly through proton pump Vacuolar ATPases (V-ATPase) activity (Banerjee & Kane, 2020; Kissing et al., 2015). This acidification is critical to halt growth of pathogenic microorganisms and is necessary for diverse microbicidal functions of the phagosome that depend on H^+^ concentrations, such as activation of degradative enzymes and production of reactive oxygen species (ROS) (Dragotakes et al., 2020; Westman & Grinstein, 2021).

Many pathogenic bacteria have developed various strategies to hijack the host machinery, avoid phagosome maturation, and produce a specific compartment where intracellular bacteria can survive and replicate (Kellermann et al., 2021; Omotade & Roy, 2019). *Brucella abortus, Legionella pneumophila* and *Chlamydia trachomatis* escape early from the endocytic pathway and produce unique vacuoles that contain endoplasmic reticulum and golgi membrane markers (Celli, 2019; Gitsels et al., 2019; Steiner et al., 2018). *Mycobacterium tuberculosis* and *Salmonella* block phagosome maturation and arrest the bacterial-containing vacuole at an early stage (Brumell, 2004; Queval et al., 2017; Steele-Mortimer, 2008). Alternatively, *Coxiella burnetii* interacts with the endocytic pathway to develop a replication-permissive spacious vacuole that is similar to a phagolysosome (J. Qiu & Luo, 2017; van Schaik et al., 2013). *A. baumannii*, on the other hand, has classically been considered an extracellular pathogen. Recent reports, however, have demonstrated that *A. baumannii* clinical isolates, but not “domesticated lab strains”, can replicate *in vitro* within *Acinetobacter* Containing Vacuoles (ACVs) in lung epithelial cells and macrophages (Rubio et al., 2022; Sycz and Di Venanzio et al., 2021). The trafficking pathways involved in the biogenesis of these ACV remains unknown.

In this work, we demonstrate that modern clinical isolates of *A. baumannii* can survive and replicate inside AM ACVs during a murine pulmonary infection. Moreover, we characterize the intracellular trafficking of *A. baumannii* and describe key differences between the “domesticated lab strain” 19606 and the new clinical isolate 398 in J774A.1 macrophages. We additionally demonstrate that the replicative strain generates a non-degradative ACV. Finally, we provide evidence that resistance to acidic pH and secretion of ammonia, a by-product of amino acid degradation, are key factors for the *A. baumannii* intracellular survival.

## Results

### *A. baumannii* clinical isolates develop ACVs in alveolar macrophages *in vivo*

Macrophages play a critical role in host immune defense against *A. baumannii* respiratory infections (H. Qiu et al., 2012). It has been reported that macrophages can phagocytose and degrade common lab strains of *A. baumannii* such as 19606, but diverse new clinical isolates persist and/or replicate in mouse and human macrophages (Sato et al., 2019; Sycz and Di venanzio et al., 2021). We have previously demonstrated that *A. baumannii* new clinical isolates replicate in macrophages inside an *Acinetobacter* Containing Vacuole (ACV) (Sycz and Di Venanzio et al., 2021). Yet, to the best of our knowledge, there is no evidence showing that *A. baumannii* replicates inside macrophages *in vivo*. To analyze that, we evaluated the ability of two *A. baumannii* strains, the “domesticated lab strain” 19606 and the recent clinical isolate 398, to produce ACVs in macrophages during an acute lower respiratory tract infection into C57BL/6 mice. Both *A. baumannii* strains expressing GFP were each inoculated intranasally in C57BL/6 mice (∼1 × 10^8^ CFU), and after 3 and 24 hpi, the number of total and intracellular colony forming units (CFUs) present in the bronchoalveolar lavage fluid (BALF) was determined by antibiotic protection assays. Additionally, the number of total and infected AMs were analyzed using flow cytometry. Finally, the presence of ACVs in macrophages was established by confocal microscopy (Fig. 1A). CFU enumeration demonstrated similar numbers of total bacteria for both *A. baumannii* strains at 3 hpi, indicating that similar numbers of bacteria reached the lung. However, the number of intracellular bacteria was greater for 398 than for 19606 (Fig 1B, left). At 24 hpi, both the total and intracellular CFUs for 398 were significantly higher compared to 19606 (Fig. 1B right). In agreement with Qiu et al., (2012), we detected an slight increase in the number of AMs between 3 hpi and 24 hpi by flow cytometry, however at 24 hpi neutrophils were the most prevalent immune cells (Fig. S1A). The total number of AMs present in the BALF was similar between mice infected with both *A. baumannii* strains, but the number of AMs infected with 398 persisted over time, while the ones infected with 19606 were significantly more cleared at 24 hpi (Fig. 1C). Visualization of the cells present in the BALF by confocal and transmission electron microscopy (TEM) showed that only 398 was able to produce large ACVs in AMs, while the “domesticated lab strain” 19606 remained contained in single-bacterium phagosomes or was degraded by 24 hpi (Fig. 1D and S1B). Together, these experiments demonstrate that, while a lab strain is rapidly cleared, the clinical isolate 398 can infect and multiply intracellularly inside ACVs in murine AMs *in vivo*.

**Figure 1.**
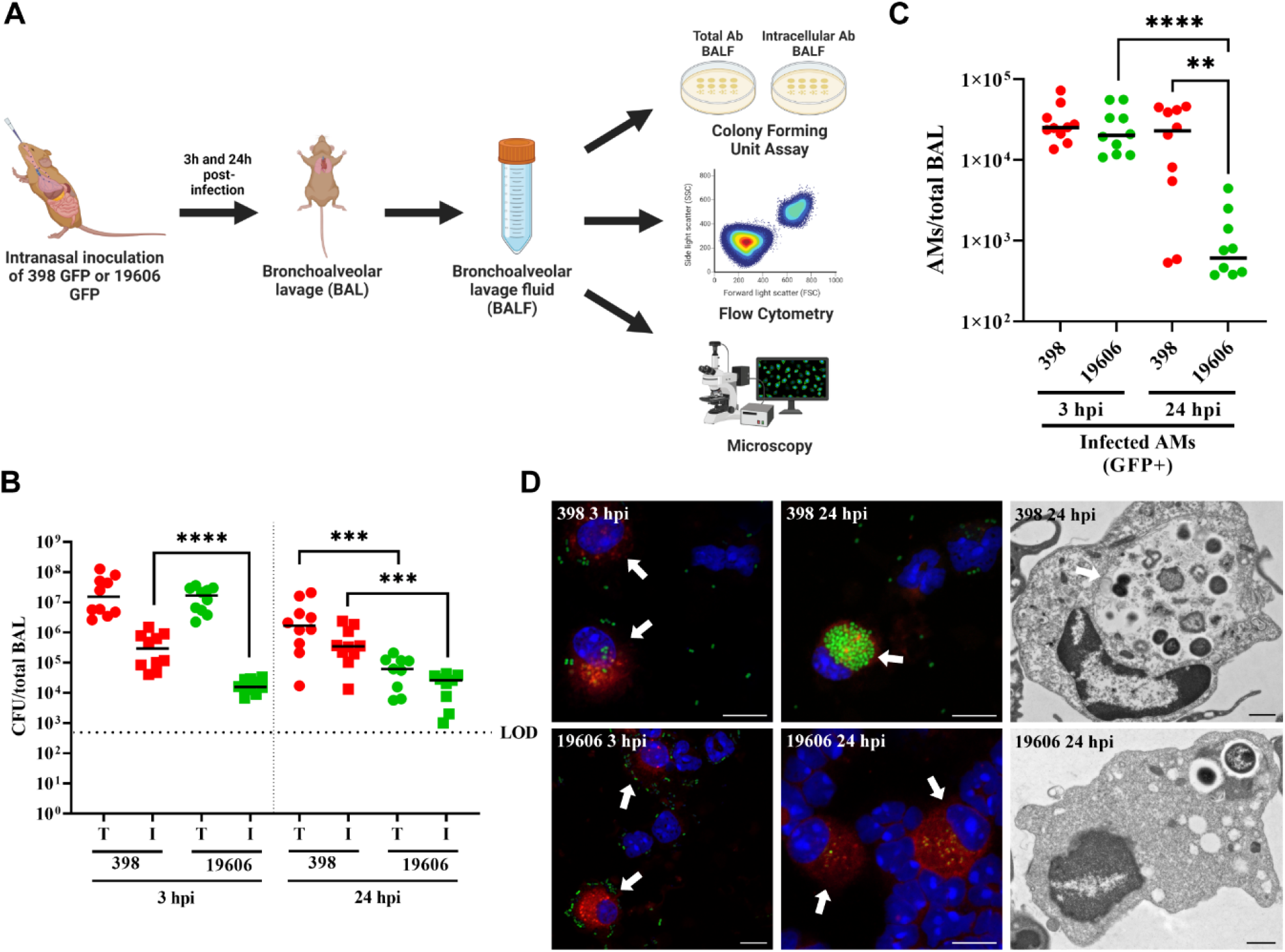
*A. baumannii* clinical isolate 398 infects AMs *in vivo* and survives inside ACVs. (A) Schematic of the pneumonia model employed in this study. (B) Mice were intranasally infected with 1 × 10^8^ CFU of *A. baumannii* strains GFP-19606 or GFP-398. At 3 or 24 hpi the total (T) or intracellular (I) CFU from bronchoalveolar lavages were determined. Symbols represents individual animals. Median values are shown as horizontal black bars. Data from two independent experiments with 5 mice per experimental group are shown. (C) Quantification of infected AMs (CD45+CD11c+SiglecF+CD11b-GFP+) obtained from BALF at 3 hpi or 24 hpi with GFP-19606 or GFP-398 strains. Symbols represent individual animals. (D) Representative confocal microscopy images of cells present in the BALF of infected mice (left and central images). Cell nuclei were stained with DAPI (blue), *A baumannii* 19606 or 398 were detected by GFP fluorescence (green), and SiglecF was immunolabeled with specific antibodies (red). Arrows indicate AMs. Scale bars: 10 μm. Representative transmission electron microscopy images with infected AMs from BALF are shown at the right. Scale bars: 1 μm. Statistical analyses were performed using the Mann–Whitney test, **p < 0,0021, ****p < 0,0001. Hpi: hours post-infection.

### The autophagic protein LC3 is absent from ACVs of a modern *A. baumannii* clinical isolate

Autophagy is a process by which eukaryotic cells remove unwanted material from the cell cytosol (Riebisch et al., 2021). Xenophagy is a highly selective type of autophagy used by the host cells to detect and eliminate intracellular pathogens (Mao & Klionsky, 2017; Sharma et al., 2018). Bacterial phagosomes can recruit the canonical autophagy marker, LC3 (microtubule-associated protein 1 light chain 3), inducing the compartment maturation and content degradation. However, the relationship between autophagy and intracellular microorganisms is complex (Grijmans et al., 2022). Some intracellular bacteria as *C. burnetii* and *Serratia marcescens* survive in an autophagic vacuole while others, such as *M. tuberculosis* and *L. pneumophila*, avoid the interaction with autophagosomes (Fedrigo et al., 2011; Mansilla Pareja et al., 2017; Omotade & Roy, 2020; Strong et al., 2022; Winchell et al., 2014). However, the relationship between the autophagic pathway and *A. baumannii* remains unclear and controversial. In epithelial cells, vacuoles of the strain AB5075 colocalize with LC3 before bacterial killing; however, ACVs of replicative strains C4 or ABC141 do not recruit the autophagic marker (Ambrosi et al., 2020; Rubio et al., 2022). In our previous work we demonstrated that, in J774A.1 macrophages, ACVs of the clinical isolate UPAB1 are LC3 negative (Sycz et al., 2021). We hypothesized that these differences could depend on the *A. baumannii* strain used in these studies. To corroborate this, the subcellular distribution of LC3 was analyzed in J774A.1 cells infected with our replicative isolate GFP-398 or the non-replicative GFP-19606. At 4 hpi, most of 19606 phagosomes were labeled with the autophagic marker (Fig. 2A bottom row, 2C and S2). On the contrary, 398 ACVs were LC3 negative up to 6 hpi (Fig. 2A-B and S2). These data suggest that while non-replicative strains such as 19606 are eliminated by autophagy, replicative strains like 398 avoid the interaction with the autophagic pathway.

**Figure 2.**
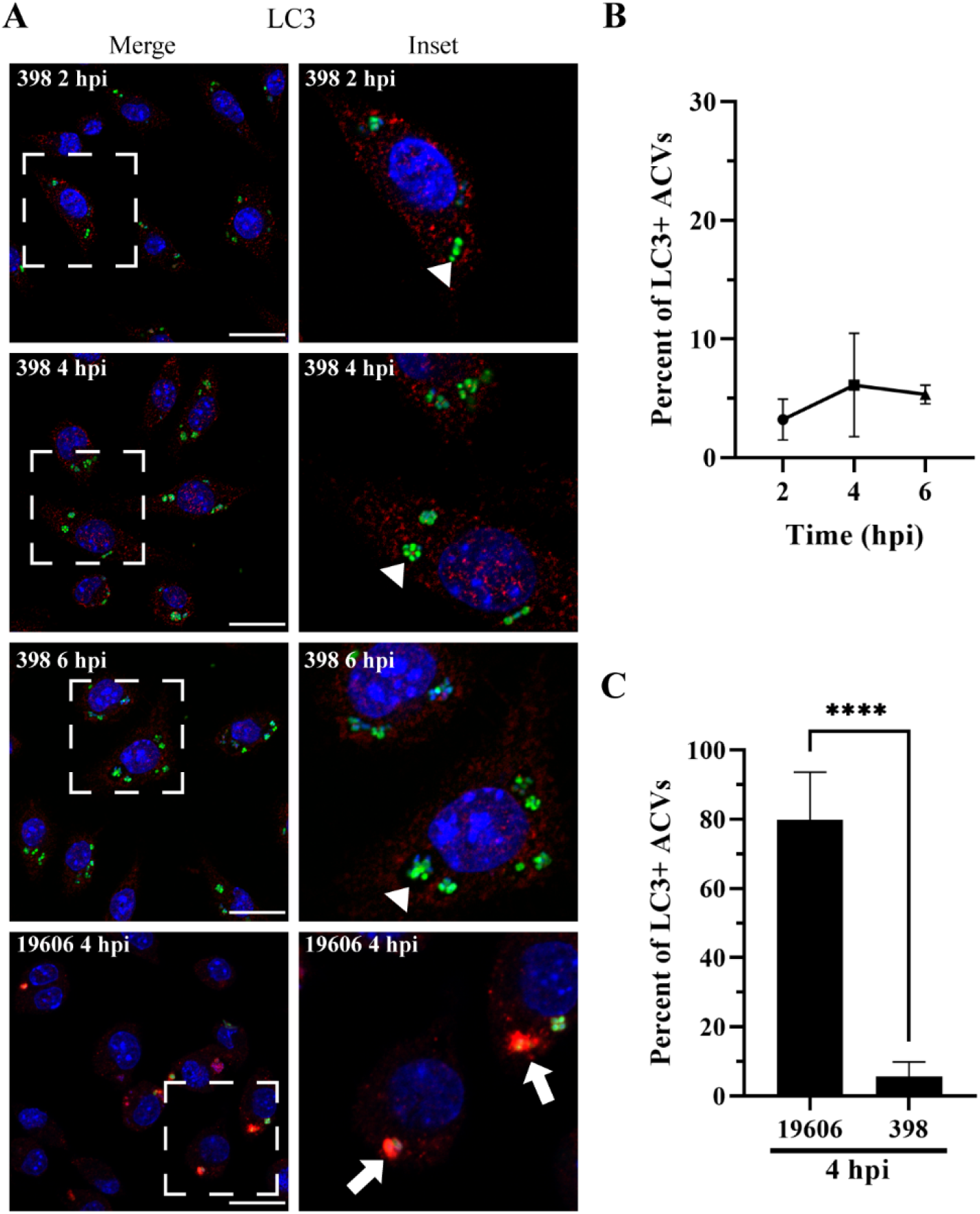
398 ACV does not colocalize with the autophagic marker LC3. (A) Representative confocal microscopy micrographs of J774A.1 cells infected with *A. baumannii* strains GFP-398 or GFP-19606 at the different times post-infection (pi). Cell nuclei were stained with DAPI (blue), bacteria overexpress GFP (green) and LC3 was immunostained with a specific antibody (red). Scale bars: 20 μm. Insets (40 μm) are a higher magnification of the area indicated with a white box in the corresponding image. LC3-positive ACVs are indicated with arrows and LC3–negative ACVs are indicated with arrowheads. (B) Quantification of LC3 positive ACVs of the clinical isolate 398 at the different times pi. (C) Comparison of the percent of LC3 positive ACVs of 398 and 19606 at 4 hpi. At least 200 infected cells were analyzed per strain per time point. The results are expressed as mean ± standard error of the mean (SEM) of three independent experiments. Statistical analyses were performed using Welch’s t-test, **** < 0.0001.

### ACVs maturate to a late phagosome

Nascent phagosomes interact with different compartments of the endocytic pathway to ultimately fuse with lysosomes and degrade their internal content. This sequential process is called “maturation” and is characterized by the recruitment of specific molecular markers to the phagosomal membrane that guide their traffic inside the cell (H.-J. Lee et al., 2020). However, several bacterial pathogens have evolved to manipulate the host cell and survive intracellularly (Kellermann et al., 2021; Omotade & Roy, 2019). To characterize the biogenesis of the ACVs, we utilized the early or late endosomal markers EEA1 (Early Endosomal Antigen 1) and LAMP1 (Lysosomal-associated membrane protein 1), respectively. Approximately, 80% of the ACVs of *A. baumannii* clinical isolate 398 colocalized with EEA1 as early as 15 min post infection (mpi), however, the percentage of colocalization of this marker decreased over time. At 60 mpi less than 5% of the ACVs were EEA1 positive (Fig. 3A,C and S3). When we analyzed the distribution of the late endosomal marker LAMP1, most of the ACVs were decorated with this marker as early as 60 mpi (Fig. 3B-C and S4A). Similar results were obtained with 19606 for LAMP1 endosomal markers (S4B-C). This data demonstrates that in macrophages the phagosome produced by both strains follow a canonical maturation, however then ACVs of replicative strains diverge towards a different pathway avoiding the interaction with autophagosomes.

**Figure 3.**
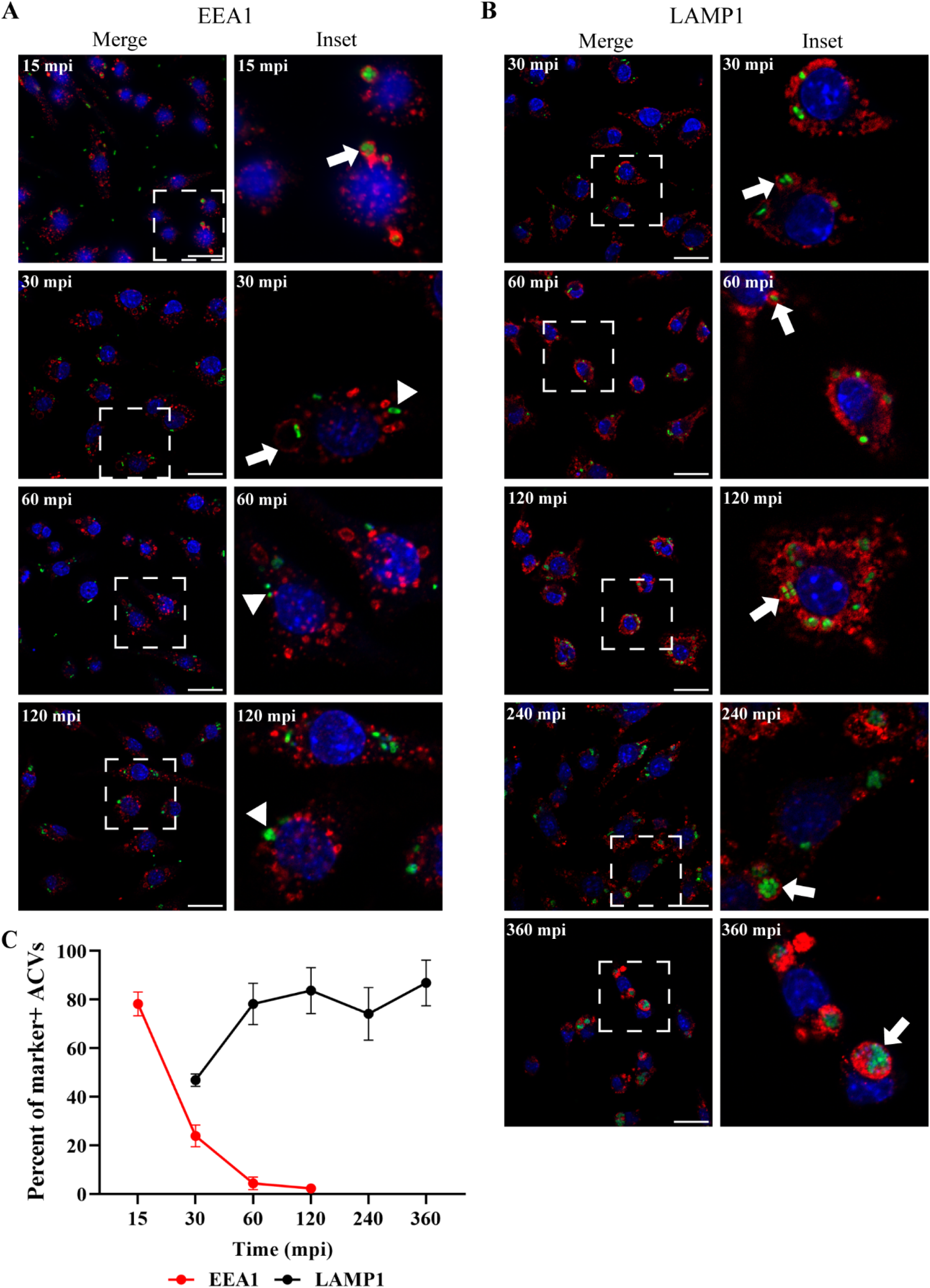
The ACV interacts with the endocytic pathway. J774A.1 cells were infected with *A. baumannii* GFP-398, and after the indicated times pi the cells were fixed and processed for confocal microscopy. The samples were stained to detect cell nuclei (blue), GFP-bacteria (green) and (A) EEA1 (red) or (B) LAMP1 (red). Bars: 20 μm. Insets (40 μm) are a higher magnification of the region indicated with a white box in the corresponding image. The presence of EEA1 or LAMP1 markers in the ACV is indicated by arrows while marker negative ACVs are denoted by arrowheads. (C) The percent of EEA1 or LAMP1 positive ACVs was determined at different times pi. At least 200 infected cells were analyzed per indicated time point. The results are expressed as mean ± SEM of three independent experiments. Mpi: minutes post-infection.

### The ACV of *A. baumannii* strain 398 is a non-degradative compartment

The final step of phagosomal maturation is the fusion with lysosomes to form the phagolysosomal compartment in which the degradation of the internal content takes place (Nguyen & Yates, 2021). To determine if the ACV is a degradative compartment, we used the fluorogenic probe DQ-BSA Green, a self-quenched substrate that fluoresces green when it is degraded by acidic proteases in the phagolysosome. The lysosomes of both uninfected and infected macrophages exhibited green fluorescence, indicating that lysosomal biogenesis and functions were not impaired overall (Fig. S5A-B). More than 50% of 19606 phagosomes were degradative compartments (DQ-BSA positives) (Fig. 4A,D and S5B bottom row). In contrast, more than 80% of 398 ACVs did not colocalize with the fluorogenic probe (Fig. 4A-C and S5B). Similarly, the median fluorescence intensity (MFI) profiles of representative ACVs show that only phagosomes containing 19606 exhibit a high green signal corresponding to the degraded DQ-BSA marker (Fig. 4B). These data indicate that the 398 ACVs are non-degradative, suggesting that 398 can manipulate the canonical maturation of the phagosome.

**Figure 4.**
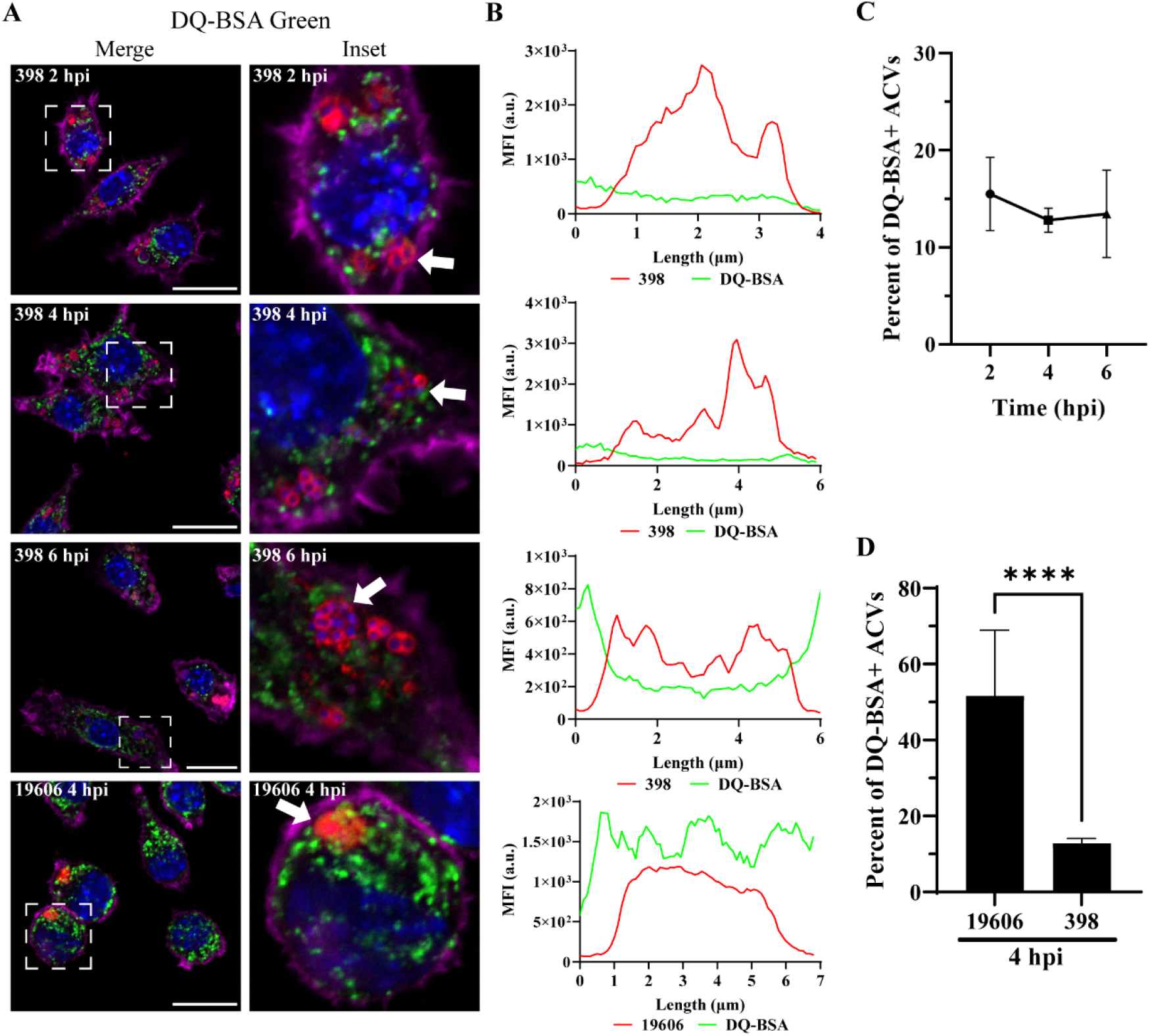
*A. baumannii* 398, but not 19606, resides in a non-degradative ACV. (A) J774A.1 cells infected with *A. baumannii* 398 or 19606 were incubated with DQ-BSA and then fixed at the indicated time points. The samples were stained to detect cell nuclei (blue), DQ-BSA (green), *A. baumannii* (red) and actin (pink). Bars: 20 μm. Insets (20 μm) are a higher magnification of the area denoted in the corresponding image with a white box. (B) Fluorescent intensity plots of representative ACVs (indicated with arrows). (C) Quantification of DQ-BSA positive ACVs of the clinical isolate 398 at the different times pi. (D) Comparison of the percentage of DQ-BSA positive ACVs of 398 and 19606 at 4 hpi. Statistical analyses were performed using the Welch’s t-test, **** < 0.0001. At least 200 infected cells were analyzed per strain, per time point. Results are expressed as mean ± SEM of three independent experiments.

### Luminal pH of the ACV increases during infection

The two main characteristics of phagolysosomes are an acidic pH and the presence of hydrolytic enzymes (Rosales & Uribe-Querol, 2017). Phagosome acidification is mainly achieved by V-ATPases that pump H^+^ into the luminal space of this compartment. To test the effect of vacuolar acidification on *A. baumannii* intracellular replication, we used bafilomycin A1, a specific V-ATPase inhibitor. While strain 19606 was degraded under normal conditions, in the presence of bafilomycin A1, 19606 was able to replicate (Fig. 5A). Moreover, large ACVs with many GFP-19606 bacteria were observed by confocal microscopy when incubated with bafilomycin A1 (Fig. 5B). These data suggests that the luminal pH of the ACV is key factor in thwarting intracellular replication of *A. baumannii*.

**Figure 5.**
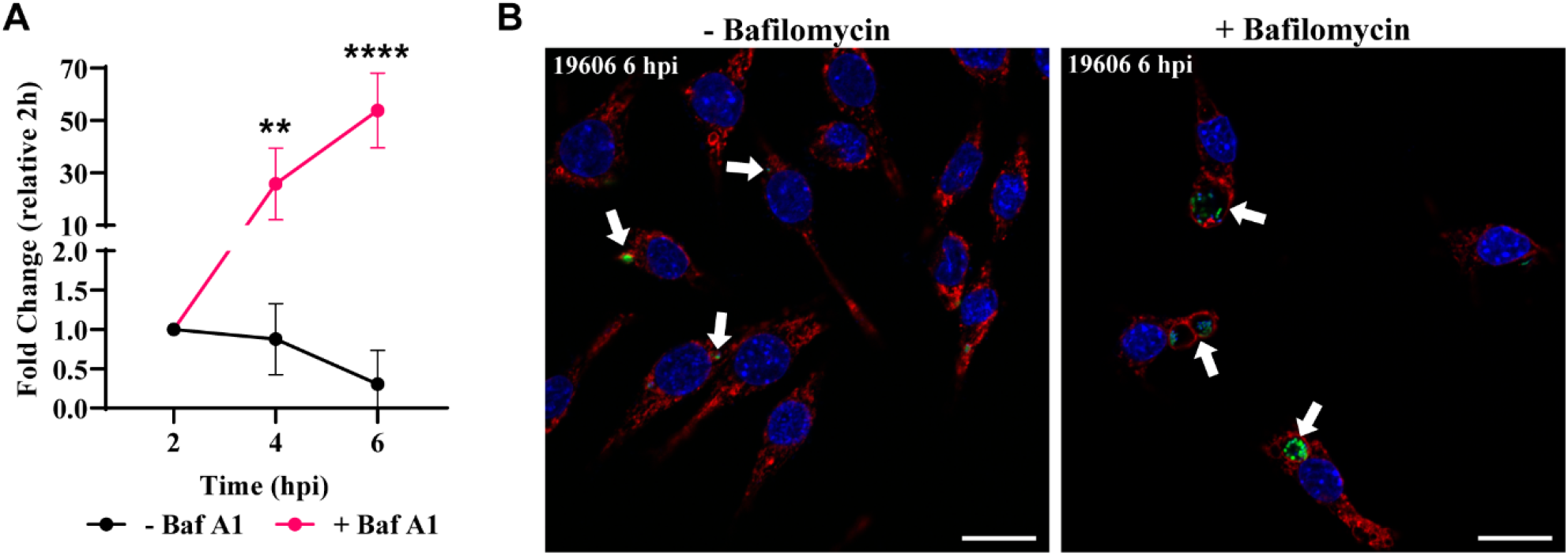
Bafilomycin A1 treatment allows 19606 to replicate in macrophages. (A) J774A.1 macrophages were infected with GFP-19606 and treated with the proton pump V-ATPase inhibitor bafilomycin A1. Total numbers of intracellular CFU were determined at different times pi in treated and non-treated cells. Statistical analyses were performed by two-way ANOVA-test, ** < 0.0021, **** < 0.0001. (B) Representative images of cells infected with GFP-19606 (green) and incubated with or without bafilomycin A1 at 6 hpi are shown. Cell nuclei were stained with DAPI (blue) and LAMP1 with specific antibody (red). Bars: 20 μm.

The ability of 398 to survive and replicate within ACVs prompted us to analyze their pH. LysoSensor is a dye that is internalized in the cell, and its fluorescence intensity is dependent on the pH of the compartment in which it resides (increased brightness at low pH). Live confocal microscopy demonstrated the presence of LysoSensor (green) in the ACVs (red) early during infection (Fig. 6A, 2 and 4 hpi white arrow), indicating that most of the vacuoles were initially acidic. However, as the infection progresses and the bacteria replicate inside the vacuole, the lumen of the ACV decreases in green signal intensity (Fig. 6A, 6 and 24 hpi white arrow), revealing a remarkable increase in the luminal pH of this compartment (Fig. 6A-B). Moreover, the number of ACVs that colocalize with LysoSensor decreased significantly over the time (Fig. 6C). These results demonstrate that although 398 vacuoles are initially acidic, this strain can actively increase the ACV’s luminal pH.

**Figure 6.**
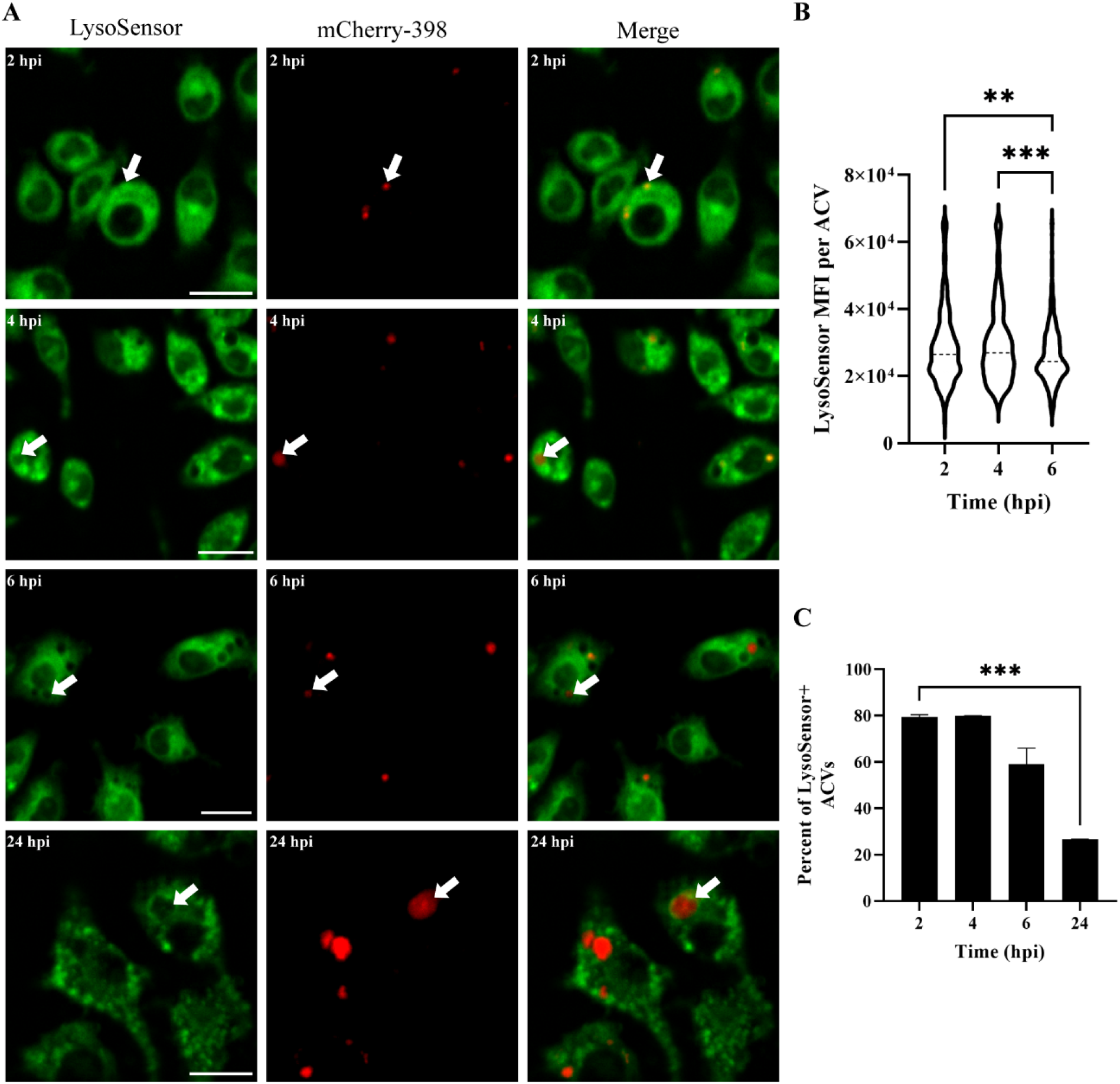
398 increases the luminal pH of the ACV during infection. (A) J774A.1 cells infected with mCherry-398 (red) were incubated with LysoSensor (green) 15 minutes before the indicated time points. Samples were analyzed by *in vivo* confocal microscopy. Bars: 20 μm. (B) Analysis of the Mean Fluorescence Intensity (MFI) signal of LysoSensor per ACV at different times pi. (C) Percentage of ACVs that colocalize with LysoSensor at 2, 4, 6 and 24 hpi. Statistical analyses were performed using one way ANOVA-test, ** < 0.0021, *** < 0.0002. At least 200 infected cells were analyzed per indicated time point. Results are expressed as mean ± SEM of three independent experiments.

### Replicative clinical isolate 398 grow better than the non-replicative strain at acidic pH

Our data suggests that the ACV pH is key in controlling *A. baumannii* replication and that the clinical isolate 398 actively increases the pH of the ACV after initially surviving acidic conditions. Previous work observed that *Acinetobacter calcoaceticus* clinical isolates are able to survive at acidic conditions (pH 4 to 6), but growth was significantly reduced compared to neutral pH (Glover et al., 2022). Also, *A. baumannii* was found to be more tolerant to pH stress than other *Acinetobacter* species such as *A. nosocomialis, A. pittii* and *A. calcoaceticus* (Peleg et al., 2012). We hypothesized that replicative strains of *A. baumannii* are more tolerant to acid stress than non-replicative bacteria. To analyze the relationship between intracellular replication and growth fitness at acidic pH, we employed the replicative strain 398 and the non-replicative strains 19606 (Fig. 7A). We tested the ability of these strains to grow in buffered LB broth at pH 5, 6, or 7, as previously described. Although the two strains grew similarly at neutral pH (Fig. S6B), 19606 had a lower growth rate than 398 at pH 5 (Fig. 7B). In non-buffered LB (pH ≈5), tested strains alkalinized the culture media to pH≈8 after growth for 18 h. However, 398 required significantly less time to neutralize the media than the non-replicative strain. For example, 398 reaches a pH of ≈7 at 6 h, while 19606 requires ≈10 h to reach the same pH (Fig. 7C). *A. baumannii* is unable to grow using most sugars individually (Cook & Fewson, 1973), instead, it heavily relies on amino acids. To feed the TCA cycle, the α-amino groups of the amino acids are removed with the concomitant production of ammonia, which is secreted to the media, resulting in an increase of the pH (Actis et al., 1999). We measured the secretion of ammonia by the two strains and found that 398 produces significantly higher amounts of ammonia than 19606 (Fig. 7D). Employing additional strains, we observed that intracellular replication correlates with the ability of the strains to withstand acidic pH and to produce ammonia (Fig S7). Together, our results suggest that both, growth at low pH and ammonia production, are key players for *A. baumannii* replication in the ACV.

**Figure 7.**
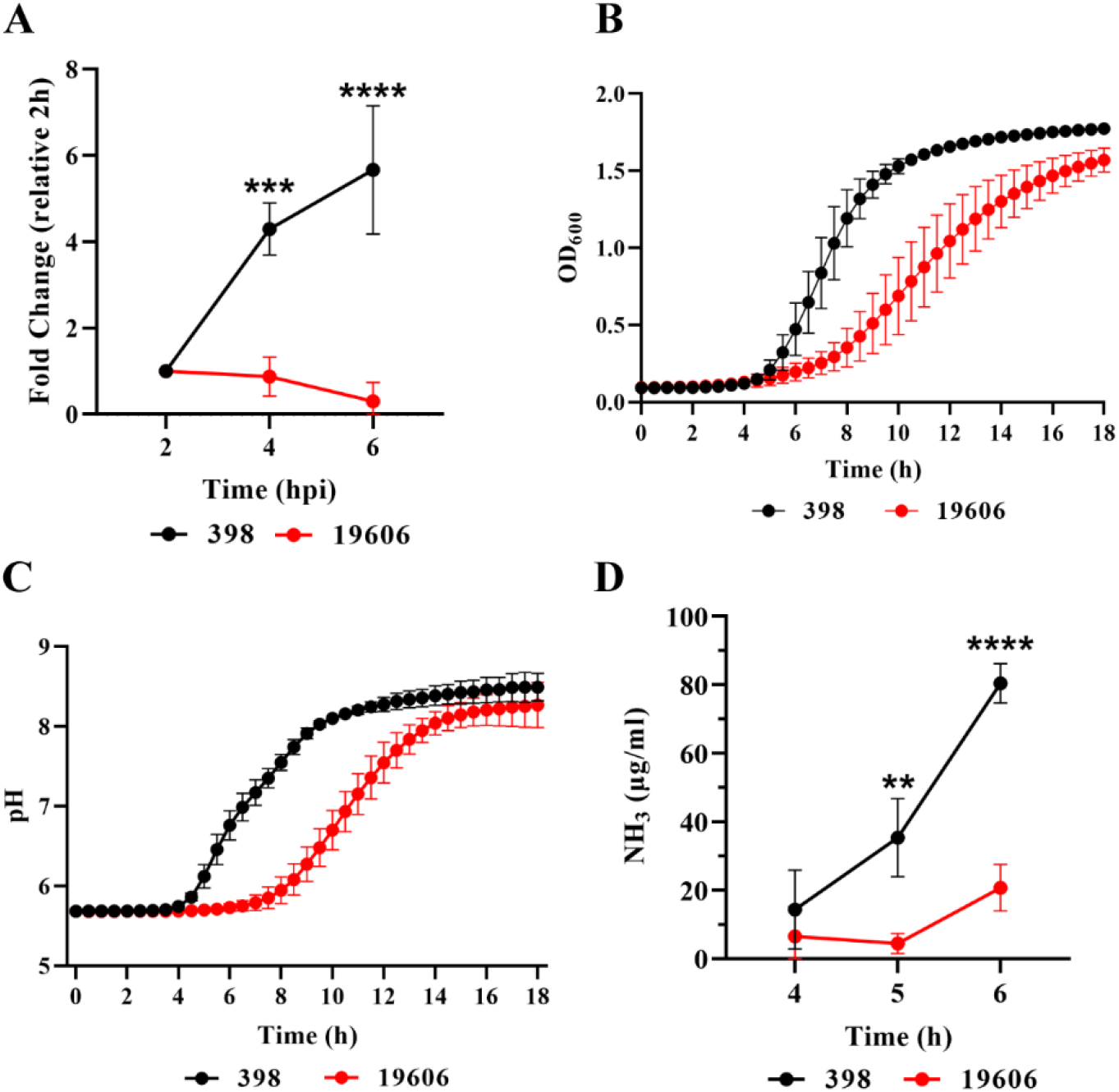
Intracellular replication correlates with the ability to grow at acidic pH. (A) The intracellular replication of *A. baumannii* strains 398 and 19606 in J774A.1 macrophages was determined by antibiotic protection assays. (B) Growth curves of 398 and 19606 in buffered LB pH 5 as measured by OD_600_. (C) Changes in culture pH during *A. baumannii* growth, determined by phenol red absorbance at 560 nm. (D) Concentration of ammonia in LB cultures of 398 and 19606 strains at 4, 5 and 6 h post-inoculation. Results are expressed as mean ± SEM of three independent experiments.

## Discussion

Here, we demonstrate that classic *A. baumannii* lab strains, like 19606, and recent clinical isolates, such as 398, behave differently, both *in vivo* and *in vitro*. While 398 was able to replicate in large ACVs within AMs during murine respiratory infection, 19606 was quickly cleared by these cells. Figure 8 summarizes our findings related to the intracellular behavior of both replicative and non-replicative strains *in vitro*. Initially, both strains follow the canonical endocytic pathway, as determined by the sequential colocalization of EEA1 and LAMP1 markers with phagosomes. 19606 phagosomes continue with their maturation and fuse with autophagosomes and lysosomes to ultimately be degraded by the macrophage. However, our data show that 398 ACVs avoid the autophagic and degradative pathways allowing the bacteria to survive and replicate inside macrophages. Moreover, 398 reverts the initial luminal acidification of the vacuole, most likely via production of ammonia, a by-product of amino acid degradation. The process generates a non-degradative niche that allows replication of *A. baumannii* inside the ACV.

**Figure 8.**
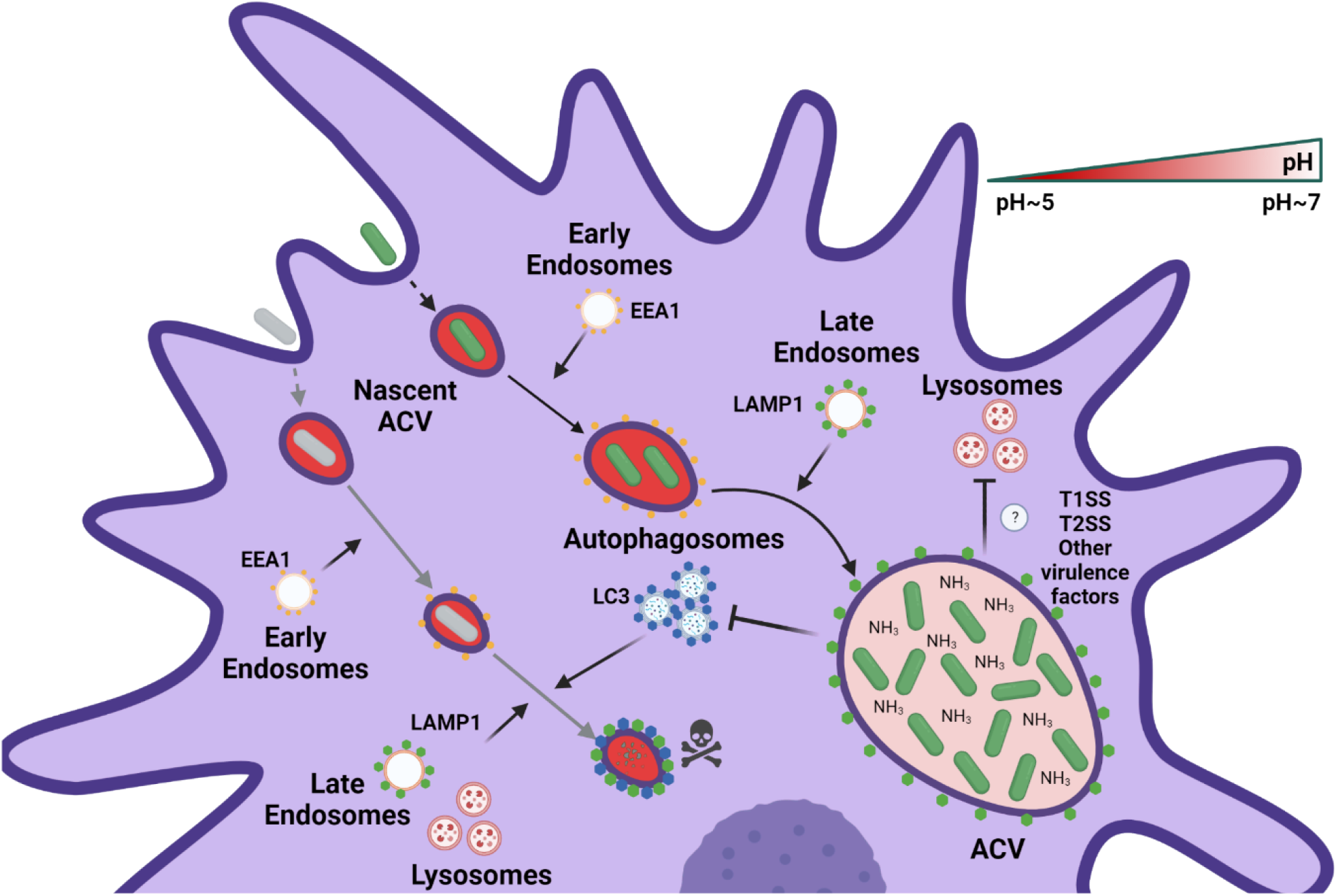
Proposed model of the intracellular lifestyle of *A. baumannii* in macrophages. Replicative strains of *A. baumannii* (green bacterium) interact with the endocytic pathway sequentially acquiring EEA1 and LAMP1 markers in the ACV. However, during its maturation, the ACV does not interact with autophagosomes or lysosomes of the host cell. Moreover, the replication of intracellular bacteria produces ammonia which neutralizes the luminal microenvironment of the ACV. On the other hand, non-replicative strains of *A. baumannii* (gray bacterium) are phagocytized by macrophages and reside in a nascent phagosome. This compartment maturates to a late phagosome, like the replicative ACV, by interactions with the endocytic pathway. Nevertheless, the final fate of the non-replicative strains of *A. baumannii* is the degradation in an autophagolysosome.

Several reports have established the critical role of AMs during *A. baumannii* respiratory infections (Gu et al., 2021; H. H. Lee et al., 2020; H. Qiu et al., 2012; Tsay et al., 2021). Depletion experiments demonstrated that AMs are necessary to control tissue damage, bacterial sepsis and severity of the infection in an *A. baumannii* pneumonia models (H. H. Lee et al., 2020). Recent reports showed that *A. baumannii* modern clinical isolates are able to survive and/or replicate intracellularly, *in vitro*, in macrophages (Sato et al., 2019; Sycz and Di Venanzio et al., 2021) and lung epithelial cells (Rubio et al., 2022). To the best of our knowledge, this is the first demonstration of the presence of large vacuoles containing *A. baumannii* within AMs in a mammalian pneumonia model. A similar behavior has been described for other pathogens, such as *M. tuberculosis, Staphylococcus aureus, and L. pneumophila*, in which bacteria use alveolar macrophages as a reservoir to persist and replicate within the host (Cohen et al., 2018; Huang et al., 2018; Khweek & Amer, 2010; Lacoma et al., 2017). Many extracellular and intracellular bacteria egressing from infected macrophages may be eliminated by neutrophils, which are the predominant cell population in mice BALF at 24 hpi (≈95%) (Fig S1A). However, *A. baumannii* predominantly infects immunocompromised patients, many of which lack neutrophils or normal neutrophil responses. For example, recent work showed that cyclophosphamide-treated mice were not able to increase the number of neutrophils in the lung after *A. baumannii* infection, leading to higher mortality (Liu et al., 2020). Therefore, replication in AMs could be a key factor for the persistence of *A. baumannii* within the infected hosts.

We demonstrated that the intracellular compartment produced by replicative *A. baumannii* strains such as 398 was clearly different than the classical phagosome formed by non-replicative strains such as 19606. Avoiding fusion with autophagosomes is critical for some intracellular bacteria to escape from degradation. The interaction between *A. baumannii* and autophagy has been studied in different host cells, but the conclusions were disparate. In epithelial cells, non-replicative *A. baumannii* strains induce autophagy, leading in some cases to persistence (Parra-Millán et al., 2018; Spiering & Hodgson, 2011), or degradation (Ambrosi et al., 2020; Wang et al., 2016). In THP-1 monocytes or MH-S macrophages, induction of autophagy after infection with *A. baumannii* strain 98-37-09 leads to bacterial killing by the host cell (Wang et al., 2016). Moreover, LC3 colocalized with *A. baumannii* strains AB5075 and 98-37-09 internalized by epithelial cells (Ambrosi et al., 2020; Wang et al., 2016). However, our data is in agreement with recent works that show that new clinical isolates that replicate inside ACVs in epithelial cells or macrophages do not recruit LC3 and get degraded (Rubio et al., 2022; Sycz et al., 2021).

Phagosomal acidification, produced mainly by the accumulation of V-ATPases, is critical to eliminate some bacteria (Westman & Grinstein, 2021). Several intracellular pathogens have developed strategies to avoid phagosomal acidification (Wong et al., 2011; Zhao et al., 2017). Here, we observed that initially the ACV of the replicative isolate 398 is acidic; however, as the infection progresses, the strain can actively increase the internal pH of the compartment. Moreover, we determined that *A. baumannii* replicative strains can better withstand acidic conditions than non-replicative strains. This allows these strains release higher levels of ammonia, leading to a quicker neutralization of the vacuolar pH (Fig. 7 and S7). While most prominent Gram-negative intracellular pathogens use a type III or IV secretion system (Du et al., 2016; Hayek et al., 2019) to manipulate the host cell machinery, *A. baumannii* only has a Type I secretion systems (T1SS) and Type II secretion systems (T2SS) (Weber et al., 2017). Multiple reports have established that the presence of ammonia in the phagosome inhibits the fusion of this compartment with lysosomes in macrophages (Gordon et al., 1980; P D’Arcy Hart & Young, 1985; P D′ Hart & Young, 1991). Other pathogens such as *Cryptococcus neoformans* or *Helicobacter pylori* can increase the phagolysosomal pH expressing a urease to produce ammonia (Fu et al., 2018; Schwartz, 2006). *M. tuberculosis* secretes the asparaginase AnsA and avoids phagosomal acidification by the production of ammonia, enabling intracellular replication (Gouzy et al., 2014). We propose a model in which *A. baumannii* strains that resist the initial acidic stress of the phagosome are able to grow and release ammonia as a metabolic by-product, thereby neutralizing the microenvironment of the ACV. Future work will determine if *A. baumannii* employs T1SS or T2SS effectors to aid in preventing acidification of the ACV and intracellular replication.

An increasing number of studies, including this one, demonstrate differences in intracellular behavior within macrophages among *A. baumannii* isolates (Dikshit et al., 2018; Li et al., 2022; Rubio et al., 2022; Sato et al., 2019; Sycz et al., 2021). Based on this evidence, we propose to redefine *A. baumannii* as a facultative intracellular bacterium. From the results obtained in this work, it is evident that *A. baumannii* can develop ACVs in AMs *in vivo* in a murine pneumonia model and it is not a phenomenon restricted to *in vitro* observations. We recently demonstrated that the clinical isolate UPAB1 establishes small intracellular reservoirs, known as ABIR (*A. baumannii* intracellular reservoirs) in bladder epithelial cells of mice that previously cleared a urinary tract infection. We observed that these ABIRs likely act as reservoirs that can be activated upon insertion of a medical device, such a catheter, leading to a resurgent infection in mice (Hazen et al., 2023). It is tempting to speculate that intracellular *A. baumannii* in AMs may also function as reservoirs in the host. We hypothesize that this intracellular lifestyle of *A. baumannii* enhances persistence inside the host. Within AMs, *A. baumannii* is protected from the immune system and from antibiotics that do not penetrate eukaryotic cells, like aminoglycosides and polymyxins.

Here, we have demonstrated that some *A. baumannii* strains are able to manipulate the host intracellular environment to form a replicative niche. Future work will focus on the identification of additional factors that *A. baumannii* employs to survive inside these host cells, which may lead to improved therapeutic interventions to treat modern *A. baumannii* clinical isolates.

## Materials and Methods

### Bacterial strains and culture conditions

The bacterial strains employed in this work are described in table S1. Bacteria were seeded on lysogeny broth (LB) agar plates and incubated 16 h at 37 °C. Thereafter, colonies were grown in LB broth under shaking conditions (200 rpm) for 16 h at 37 °C. When appropriate, antibiotics were added to the cultures at the following concentrations: 50 μg/ml zeocin, 30 μg/ml kanamycin, and 15 μg/ml chloramphenicol.

### Growth assays

*A. baumannii* strains 19606 and 398 were cultured overnight in LB broth at 37 °C under shaking conditions. Bacterial cultures were centrifugated at 6500 rpm for 5 min and the pellets were washed with Phosphate-buffered saline (PBS, Sigma, D8662). *A. baumannii* suspensions were diluted to OD_600_ = 0.01 in 150 μl of LB buffered at pH 5 (MES 100 mM), 6 (MES 100 mM), or 7 (HEPES 100 mM) in 96-well plates and incubated 16 h at 37 °C under shaking conditions. OD_600_ values were measured at 30 min intervals using a BioTek microplate spectrophotometer. The experiment was performed in biological triplicate with three technical replicates per experiment for each strain and condition.

### pH measurements in *A. baumannii* cultures

The absorbance of phenol red solutions at 560 nm is an indicator of pH, as previously described (Held, 2018; Paye et al., 2018; Rovati et al., 2012). To analyze the pH changes in *A. baumannii* cultures during bacterial growth, indicated strains were maintained overnight in LB broth at 37 °C under shaking conditions. Bacterial cultures were washed and diluted as described for growth assays, with the following modification: non-buffered LB pH 5 with phenol red (15 μg/ml) was used. OD_600_ and OD_560_ values were measured every 30 min for 16 h using a BioTek microplate spectrophotometer. The OD_600_ absorbance values were subtracted from the absorbance at 560 nm data to exclude the effect of bacterial density. Finally, the values obtained were used to estimate culture pH by employing a standard curve of OD_560_ vs pH. The experiment was performed in triplicate with three technical replicates per each strain.

### Ammonia production

Ammonia production by *A. baumannii* strains 19606 and 398 was measured using the Ammonia Assay Kit (AA0100, Sigma) according to the manufacturer’s instructions. Briefly, the strains were cultured overnight in LB broth at 37 °C under shaking conditions. Then, bacterial cultures were centrifugated, and washed with PBS. *A. baumannii* suspensions were diluted to OD_600_ = 0.01 in 150 μl of buffered LB (MES 100 mM, pH 5) in 96 well plates and incubated 6 h at 37 °C under shaking conditions. Ammonia levels were measured, at the indicated times. The experiment was performed in biological triplicate.

### Cell culture conditions

The J774A.1 mouse macrophage cell line (ATCC TIB-67) was cultured in Dulbecco’s Modified Eagle Medium (DMEM) High Glucose (Hyclone, SH30022.01) supplemented with 10% heat-inactivated Fetal Bovine Serum (FBS, Corning) at 37°C and 5% CO_2_.

### Murine Model of *A. baumannii* pneumonia

Colonies from *A. baumannii* strains 398, GFP-398 and GFP-19606 were subcultured from LB agar plates and grown in LB broth 16 h at 37°C under shaking conditions. Overnight cultures obtained were diluted in sterile LB broth (1:100) and incubated statically for 3 h at 37 °C under shaking conditions. The bacterial inocula were prepared by centrifugation at 6500 rpm for 5 min, washed twice in PBS, and the pellet resuspended in PBS. Six-to eight-week-old female C57BL/6 mice obtained from Charles River Laboratories were intranasally inoculated with *A. baumannii* strains 398, GFP-398, or GFP-19606. Briefly, mice were anesthetized by inhalation of 4% isoflurane and were infected immediately with 50 μL of bacterial suspension (∼1 × 10^8^ Colony Forming Units [CFU]) that were gradually released with a micropipette into the nares. At 3 and 24 hours post-infection (hpi), bronchoalveolar lavage fluid (BALF) was collected. For BALF collection, mice were euthanized by CO_2_ asphyxiation and then, the lungs and trachea were exposed by surgical dissection and polyethylene tubing (0.86×1.27mm) was inserted into the trachea. Cold PBS containing 1mM EDTA was flushed through the lungs in 1ml increments to collect a total volume of 10 ml. All pneumonia studies were performed in accordance with the guidelines of the Committee for Animal Studies at Washington University School of Medicine, and we have complied with all relevant ethical regulations.

### Antibiotic protection assay

To analyze the intracellular replication of the different strains, J774A.1 cells were seeded 16 h before the experiment in 48-well plates (3×10^5^ cells/well). *A. baumannii* colonies from LB agar plates were inoculated in LB broth and grown overnight at 37°C under shaking conditions. Bacterial cultures were centrifugated 5 min at 6500 rpm and washed twice with PBS. Then, bacterial cultures were normalized to OD_600_ ≈ 1 and appropriate volumes was added to 500 μl of DMEM per well to achieve an MOI≈10. Where indicated, bafilomycin A1 (15 nM; Sigma, B1793) was added to the growth media and maintained throughout the infection. Infected cells were centrifugated 10 min at 200 x g and incubated at 37°C and 5% CO_2_. After 1 h, macrophages were washed three times with PBS and treated with DMEM supplemented with 10% FBS and colistin (50 μg/mL) to eliminate the extracellular bacteria. At the time points indicated, the cells were washed three times with PBS and lysed with 500 μl of Triton X-100 (0.05%) per well. Serial dilutions of bacterial suspensions obtained were plated on LB agar and incubated overnight at 37 °C, to determine number of colony forming units (CFUs).

To determine the number of total and intracellular bacteria present in BALF from *A. baumannii* infected mice, two 500 μl aliquots of lavage fluid were centrifugated at 6500 rpm for 10 min. The pellets were incubated for 1 h at 37 °C in PBS (total bacteria) or PBS with colistin (50 mg/mL) (intracellular bacteria). Then the cells were washed three times with PBS and lysed with 500 μl of Triton X-100 (0.05%) per aliquot. CFUs were determined by serial dilutions of the bacterial suspensions.

### Flow Cytometry

BALF samples were centrifugated at 300 x g for 5 min and incubated in 500 μl Pharm Lyse Buffer (BD Biosciences 555899) for 3 min at room temperature to lyse red blood cells. The cells were then washed with flow cytometry buffer [PBS + 0.5% Bovine Serum Albumin (BSA) (Fisher BioReagents, BP9706100) + 2 mM EDTA] and incubated at 4°C with Fc Block (BD Biosciences, 553142) for 10 min. Samples were then stained for 30 min with CD45-BV510 (Biolegend, 103138) and Ly6G-BV421 (Biolegend,127628), CD11b-Alexa700 (BD Biosciences, 557960), CD11c-APC (BD Biosciences, 550261), and Siglec-F-BV786 (BD Biosciences, 740956). Finally, cells were washed and fixed in 2% paraformaldehyde until acquisition on a Benton Dickinson (BD) LSR II Fortessa cytometer. GFP signal was acquired in the FITC channel. Total cell counts per mouse were calculated using Precision Count Beads (Biolegend, 424902) according to the manufacturer’s instructions.

### Cytospin of BALF cells

BALF samples were centrifugated at 300 x g for 5 min and the pellets were resuspended in 500 μl Pharm Lyse Buffer (BD Biosciences, 555899) for 3 min at room temperature to lyse red blood cells. The cells were resuspended in PBS and total cells and viability was determined using Trypan Blue solution (Sigma, T8154) and counted using the TC20 automated cell counter (BioRad). Samples were centrifugated at 300 x g for 5 min onto CytoPro Poly-L-Lysine Coated Microscope Slides (ELITechGroup, SS-118). The slides were air-dried overnight at 4 °C and fixed in 4% paraformaldehyde. Samples were incubated with permeabilizing and blocking solution [PBS + 0.1% saponin + 0.5% BSA (Fisher BioReagents, BP9706100) + 10% FBS (Corning)]. Cells were stained with Alexa Fluor® 647 Rat Anti-Mouse Siglec-F antibody (BD, 562680) and nuclei with 4’,6-Diamidino-2-Phenylindole, Dihydrochloride (DAPI) solution (Invitrogen, D1306) 1 h at 37°C. After staining, the samples were rinsed with washing solution [PBS + 0.1% saponin + 0.5% BSA (Fisher BioReagents, BP9706100)], then rinsed with water, and mounted with a coverslip in Invitrogen™ ProLong™ Gold Antifade Mountant (Invitrogen, P36930). Finally, the cells were analyzed by confocal microscopy.

### Immunofluorescence staining

1.3×10^5^ J774A.1 cells were plated onto glass coverslips in 24 well plates and incubated 16 to 18 h at 37 °C and 5% CO_2_. Inocula of the indicated *A. baumannii* strains were prepared by centrifugation of overnight cultures at 6500 rpm for 5 min, washing twice in PBS, and resuspension of the pellet in PBS. Bacterial suspensions were normalized to OD≈1 and an appropriate volume was used to infect the cells (MOI≈10). Afterward, the plates were centrifugated 10 min at 200 x g to enhance bacterial contact with the host cells and incubated for 1 h at 37 °C and 5% CO_2_. Cells were washed three times with PBS, and extracellular bacteria were killed by treatment of the cells with colistin (50 μg/mL) for 1 h. When necessary, cells were incubated with DQ™ Green BSA (10 μg/ml; Invitrogen, D12050) 1 h prior to the end of the infection. At the indicated time points, samples were fixed with 4% paraformaldehyde for 15 min at 37 °C and then stored in permeabilizing and blocking solution [PBS + 0.1% saponin + 0.5% BSA (Fisher BioReagents, BP9706100) + 10% FBS (Corning)]. The glass coverslips were incubated with the indicated primary antibodies produced in rabbit: anti-LC3 (Sigma, L7543), anti-EEA1 (Invitrogen, PA1-063A), anti-LAMP1 (Abcam, ab24170) or anti-*A. baumannii* at a 1:100 dilution for 1 h at 37 °C. The cells were then washed 3 times with washing solution and incubated with the indicated secondary antibody goat anti-rabbit: Alexa Fluor 647 (Invitrogen, A-21244) at a 1:250 dilution, Alexa Fluor 555 phalloidin (0.33 μM; CST, #8953) and DAPI for 1 h at 37 °C. Afterwards, samples were washed with PBS, rinsed with water, and mounted with a coverslip in Invitrogen™ ProLong™ Gold Antifade Mountant (Invitrogen, P36930). Stained samples were analyzed by confocal microscopy.

### Confocal microscopy

Infected cells were analyzed with a Zeiss LSM880 laser scanning confocal microscope (Carl Zeiss Inc.) equipped with 405nm diode, 488nm Argon, 543nm HeNe, and 633nm HeNe lasers. A Plan-Apochromat 63X (NA 1.4) DIC objective and ZEN black 2.1 SP3 software were used for image acquisition. Live images were acquired with a Zeiss spinning disk confocal microscope (Carl Zeiss Inc.) equipped with 488nm and 560nm lasers. A Plan-Apochromat 63X/1.3 Oil Ph 3 (UV) VIS-IR M27 objective and ZEN black 2.1 SP3 software were employed for image acquisition. Images were analyzed using ImageJ software (NIH, USA).

### Live imaging

5×10^5^ J774A.1 macrophages were plated in a 10 mm Glass Bottom Culture 35 mm petri dish (MATEK corporation, P35G-0-14-C) and incubated 12-16 h at 37 °C and 5% CO_2_. The next day, cells were infected with mCherry-398 at an MOI≈10 as described above. 15 minutes prior to the indicated infection time points, cells were washed twice with PBS and incubated in DMEM with LysoSensor Green DND-189 (1μM; Molecular Probes, Invitrogen, L7535). The cells were then rinsed twice in PBS and were immediately analyzed by confocal microscopy. During image acquisition, the cells were maintained at 37 °C and 5% CO_2_ in a temperature-controlled CO_2_ chamber on the microscope.

### Transmission electron microscopy

Cells from BALF were centrifuged at 300 x g for 5 minutes and incubated in 500μl Pharm Lyse Buffer (BD Biosciences 555899) for 3 minutes at room temperature to lyse red blood cells. The cells were centrifugated and resuspended in PBS. Total cells number and viability were measured using Trypan Blue solution (Sigma T8154) and counted with TC20 automated cell counter (BioRad). At least 1×10^6^ cells were fixed in 2% paraformaldehyde/2.5% glutaraldehyde (Polysciences Inc., Warrington, PA) in 100 mM sodium cacodylate buffer pH 7.2 for 1 h at room temperature and subsequently incubated to 4 °C overnight. Samples were washed in sodium cacodylate buffer at room temperature and postfixed in 1% osmium tetroxide (Polysciences Inc.) for 1 h. The cells were then rinsed in distilled water, and bloc-stained for 1 h with 1% aqueous uranyl acetate (Ted Pella Inc., Redding, CA). Subsequently, the samples were rinsed in distilled water several times, dehydrated in a graded series of ethanol, and finally embedded in Eponate 12 resin (Ted Pella Inc.). Sections of 95 nm were cut with a Leica Ultracut UCT ultramicrotome (Leica Microsystems Inc., Bannockburn, IL), stained with uranyl acetate and lead citrate, and viewed on a JEOL 1200 EX transmission electron microscope (JEOL USA Inc., Peabody, MA) equipped with an AMT 8-megapixel digital camera and AMT Image Capture Engine V602 software (Advanced Microscopy Techniques, Woburn, MA). Images were processed using ImageJ software.

### Statistical analysis

Statistical analyses were performed using GraphPad Prism 8.0 (GraphPad Software Inc., La Jolla, CA). Datasets were analyzed by Mann–Whitney test, Welch’s t-test, one way ANOVA-test or two-way ANOVA-test, as indicated.

## Acknowledgments

We thank the imaging laboratory of the Molecular Microbiology Department at Washington University in St Louis. We want to thank Wandy Beatty for her collaboration in obtaining the electron microscopy images shown in this work. We thank the members of the Feldman lab for critical reading of the manuscript. This work was supported by National Institute of Allergy and Infectious Diseases (NIAID) Grant R01AI144120.

## Supporting Information

**Figure S1.**
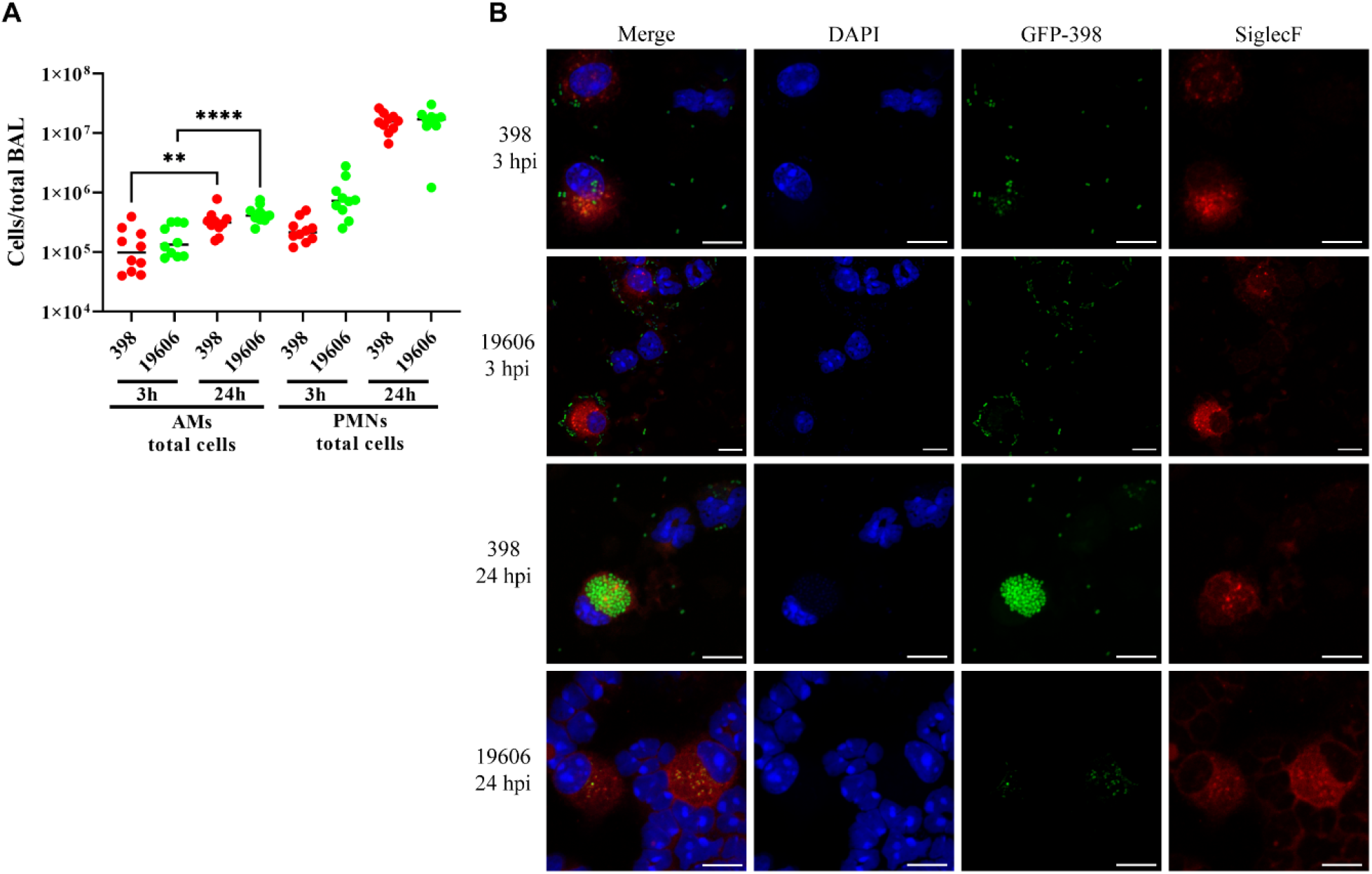
*A. baumannii* clinical isolate 398 infects AMs *in vivo* and survives inside ACVs. (A) Quantification of total AMs (CD45+CD11c+SiglecF+CD11b-) and PMNs (CD45+CD11b+Ly6G+) in the BALF of mice infected for 3 h or 24 h with GFP-398 or GFP-19606 strains. (B) Individual channels from confocal micrograph showed in the figure 1D.

**Figure S2.**
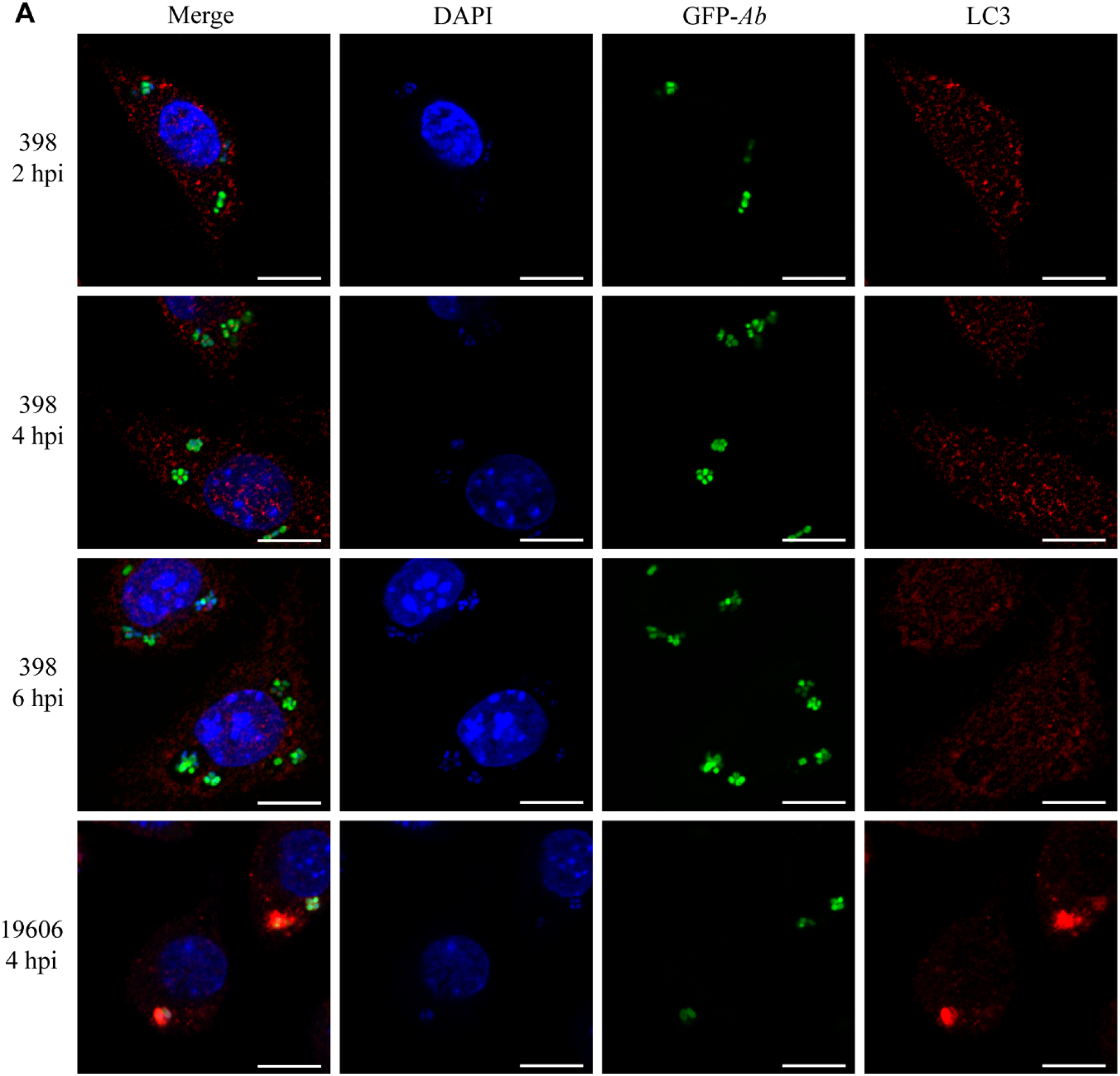
398 ACV does not colocalize with the autophagic marker LC3. (A) Single channel images of the inset micrograph shown in panel 2A. Bars: 10 μm.

**Figure S3.**
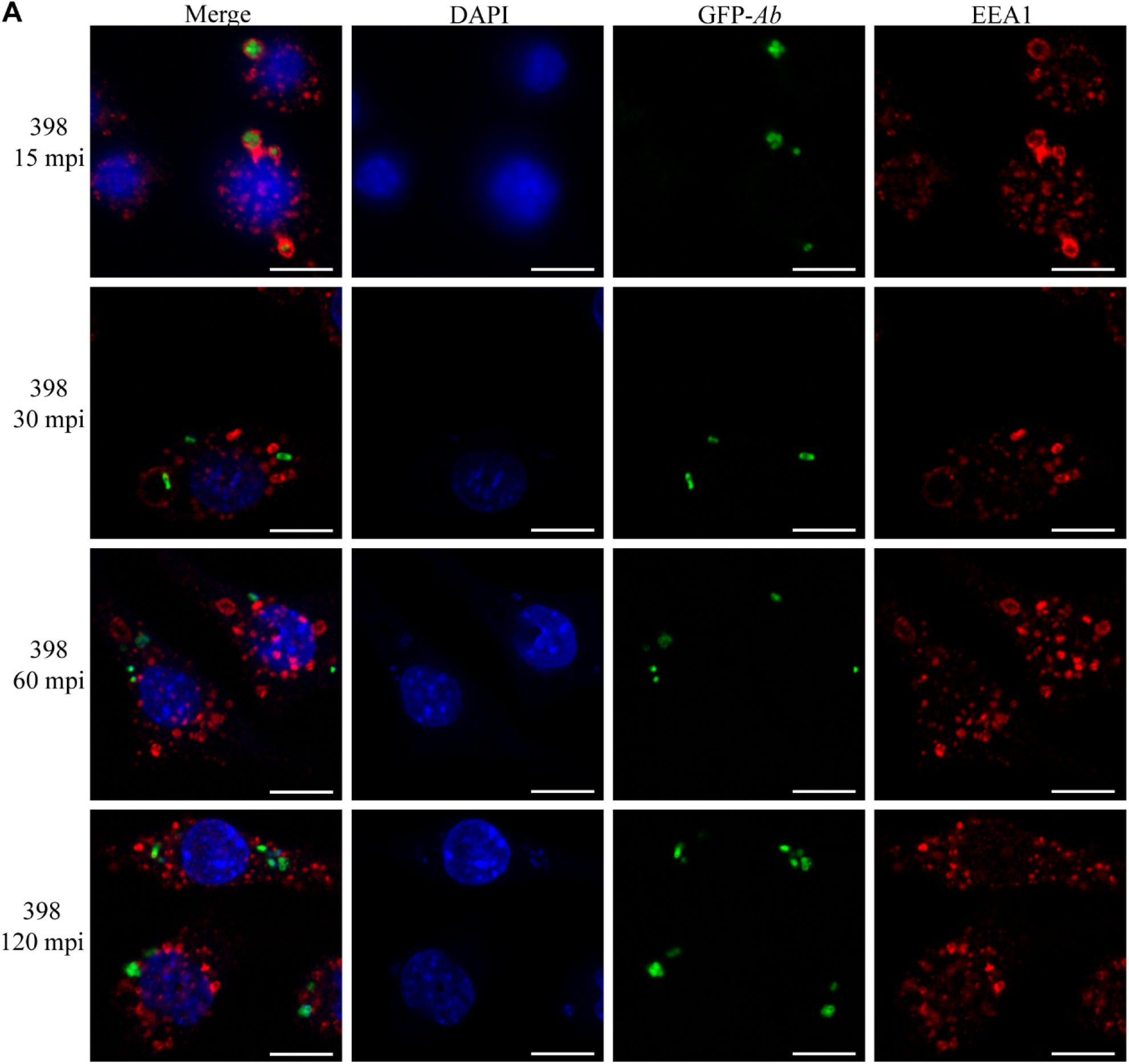
The ACV interacts with the early marker EEA1. (A) Single channel images from the inset micrograph shown in figure 3A. Bars: 10 μm.

**Figure S4.**
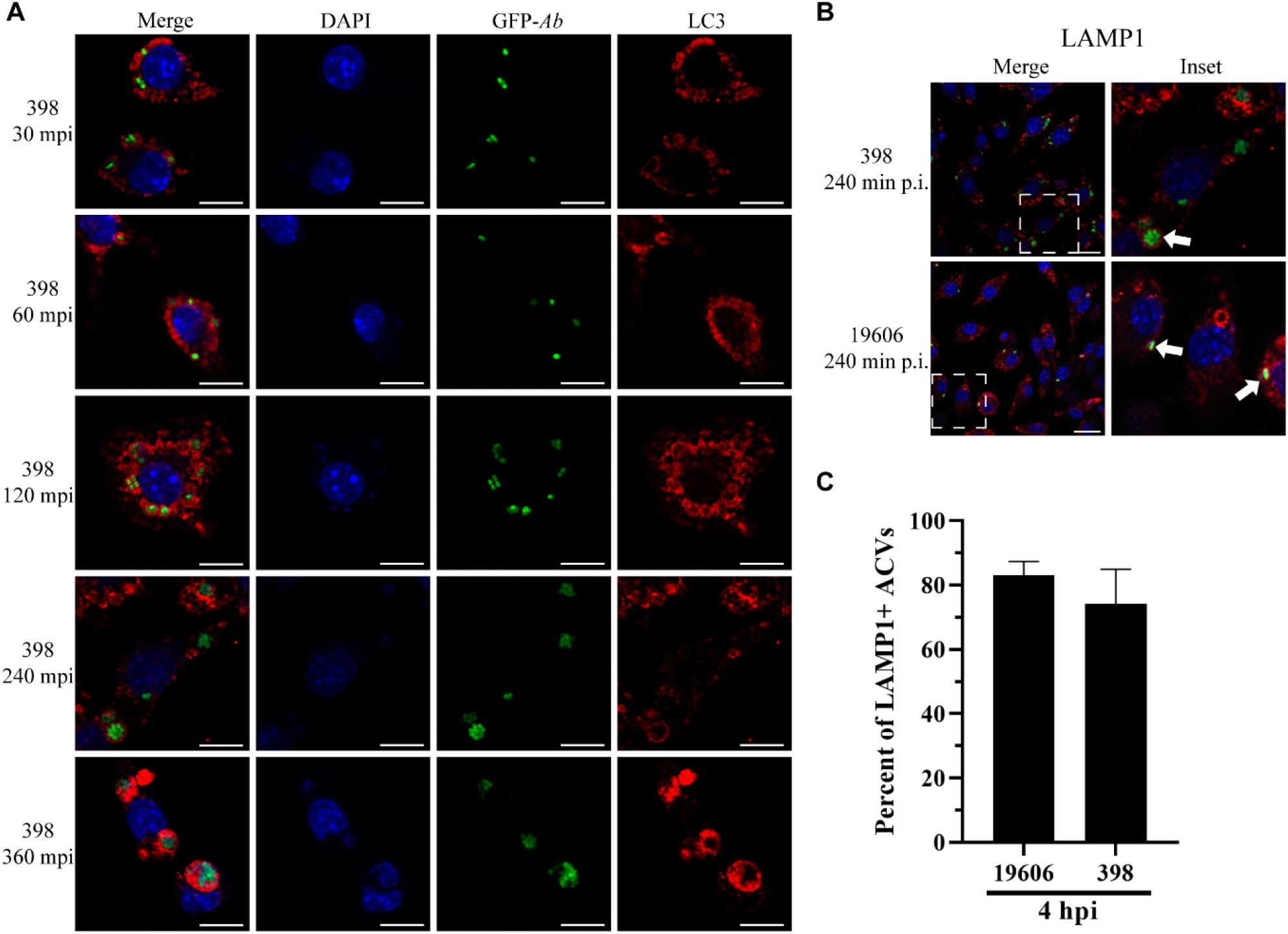
398 and 19606 ACVs colocalize with the late marker LAMP1. (A) Single channel images of the inset micrograph shown in panel 3B. Bars: 10 μm. (B) J774A.1 macrophages were infected with strains GFP-398 or GFP-19606 and fixed 4 hpi. The samples were stained to observe cell nuclei (blue), GFP-*A. baumannii* (green) and LAMP1 (red). Representative confocal images of the infections are shown. White arrows indicate ACVs that colocalize with the marker LAMP1. Bars: 20 μm. Insets (40 μm) are a higher magnification of region indicated in the corresponding image with a white box. (C) Quantification of LAMP1+ 398 or 19606 ACVs. At least 200 infected cells were analyzed. The results are expressed as means ± SEM of three independent experiments.

**Figure S5.**
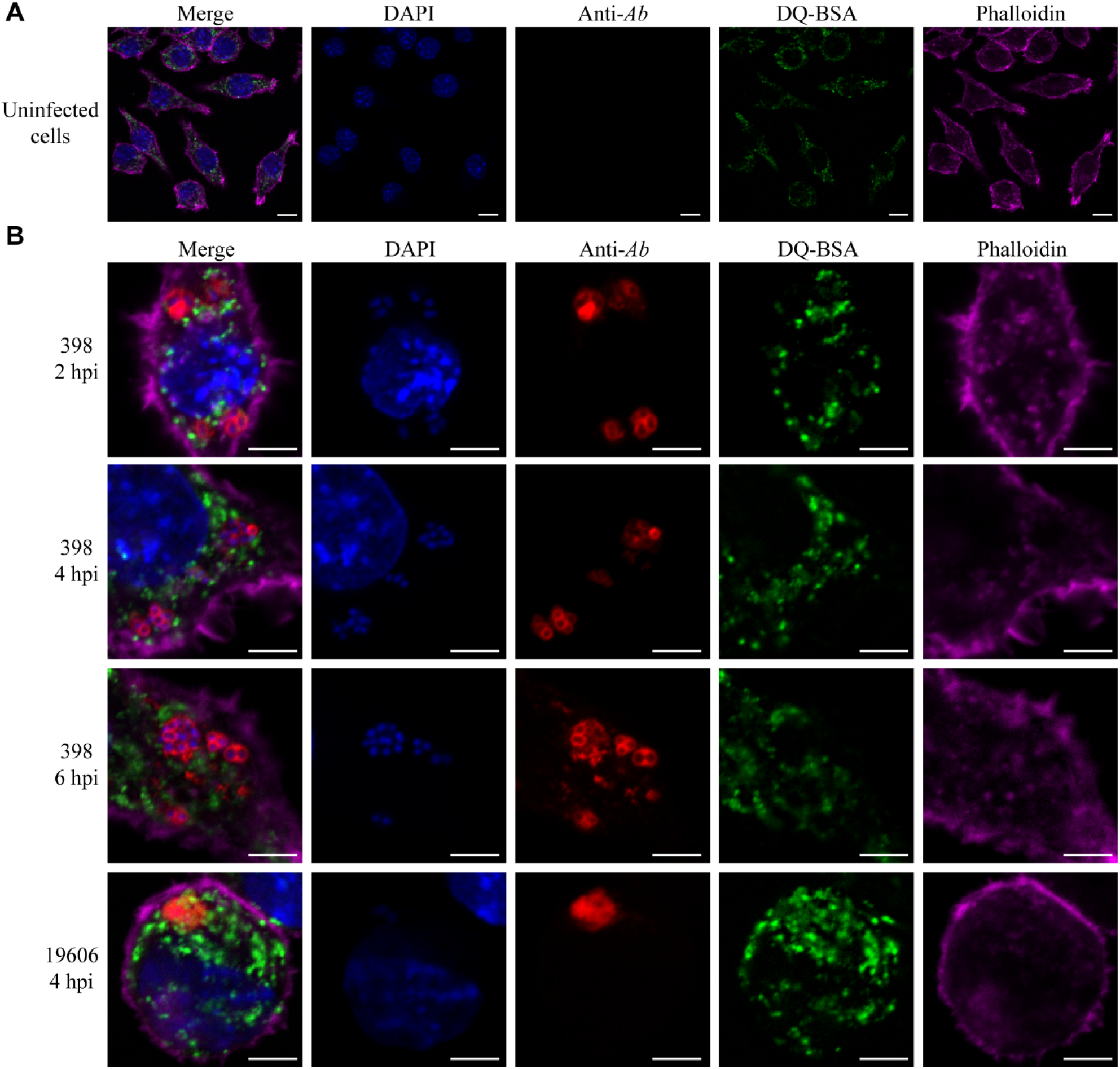
DQ-BSA green fluorescence in non-infected or *A. baumannii* infected cells. (A) Representative image of non-infected J774A.1 cells treated with DQ-BSA green. (B) Single channel images of the inset micrograph shown in figure 4A. Bars: 5 μm.

**Figure S6.**
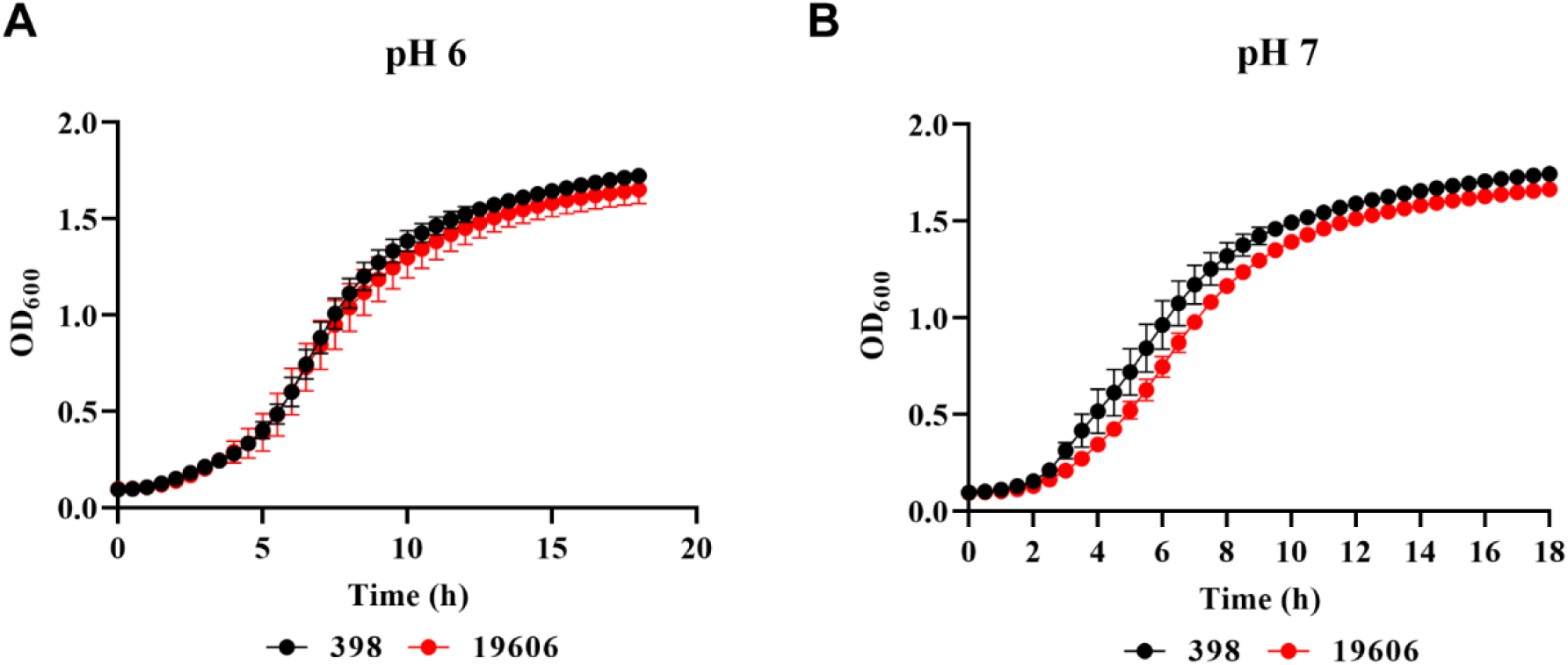
Growth of *A. baumannii* strains at different pH. Growth of 398 and 19606 strains in LB buffered at (A) pH 6 or (B) pH 7 was measured by OD_600_.

**Figure S7.**
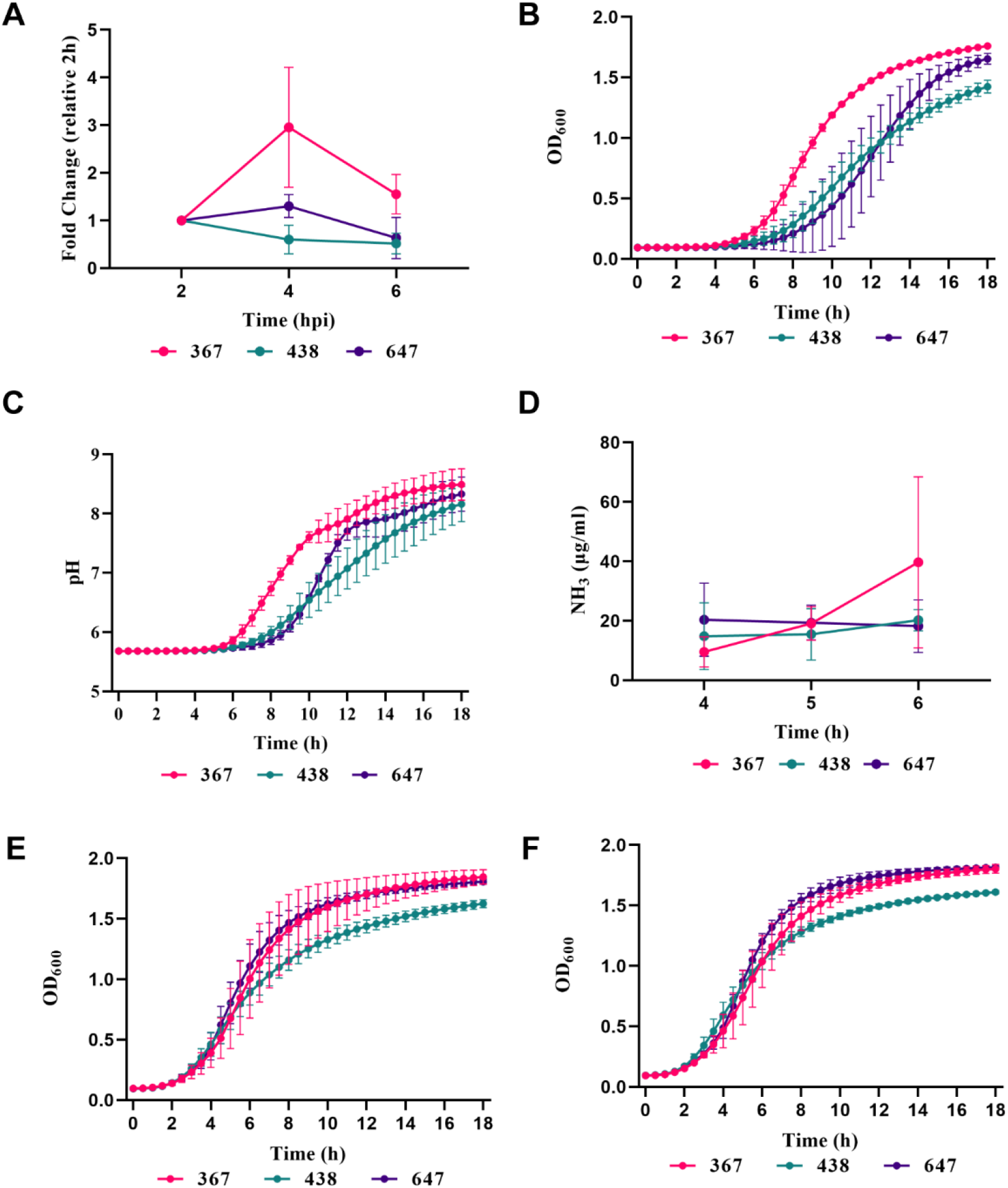
The replicative capacity of *A. baumannii* clinical isolates is related to growth fitness at acidic pH. (A) Intracellular replication of *A. baumannii* clinical isolates 367, 438 and 438 in J774A.1 macrophages determined by antibiotic protection assays. Growth of *A. baumannii* strains in LB buffered at (B) pH 5, (E) pH 6 or (F) pH 7 was measured by OD_600_. (C) Changes in culture pH during *A. baumannii* strains growth, determined by phenol red absorbance at 560 nm. (D) Concentration of ammonia in LB cultures of *A. baumannii* strains at 4, 5 and 6 h post-inoculation. Results are expressed as mean ± SEM of three independent experiments.

**Table S1.**
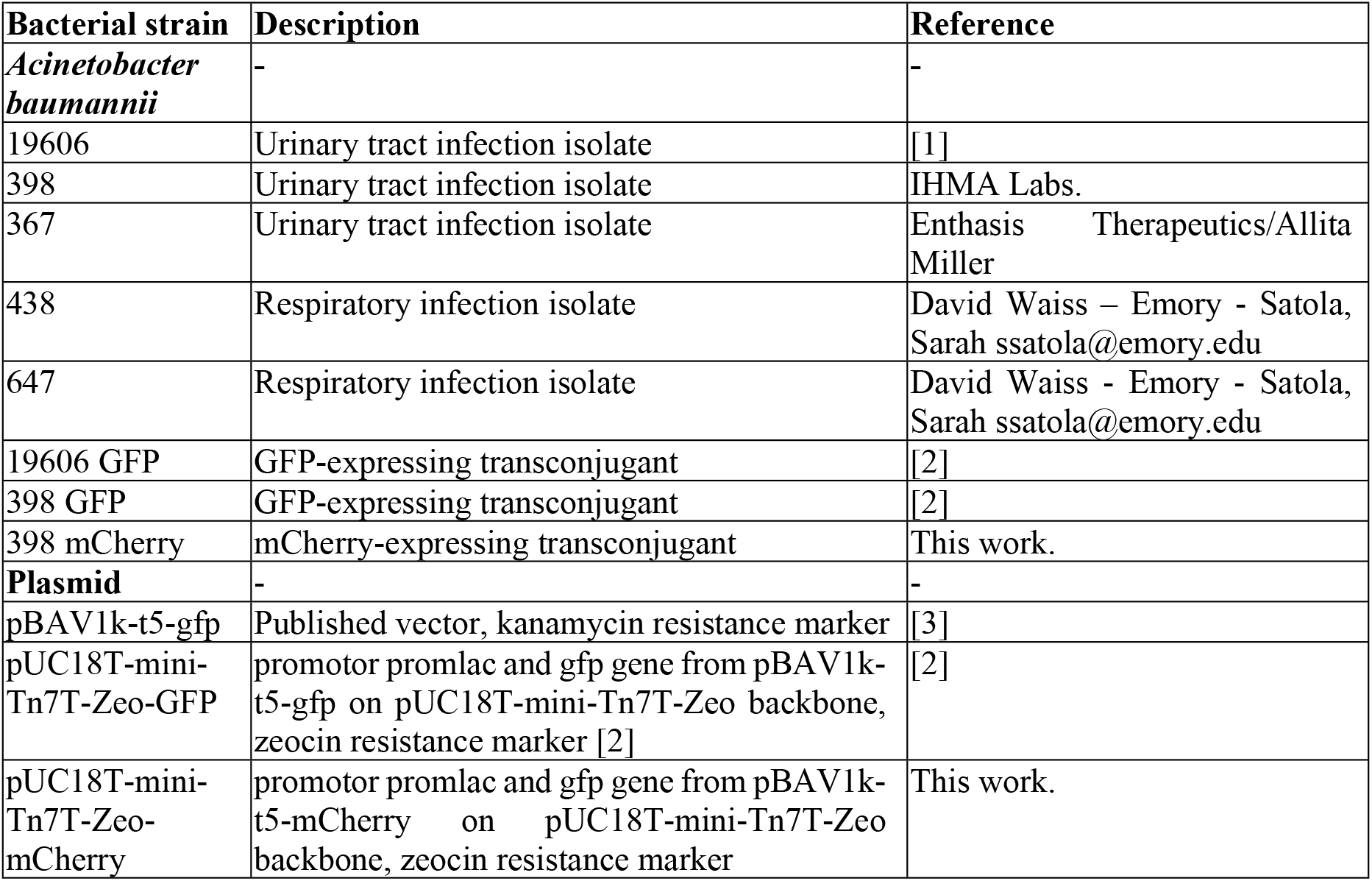
Bacterial strains and plasmids.

